# Effects of salvianolic acid A on β-amyloid mediated toxicity in *Caenorhabditis elegans* model of Alzheimer’s disease

**DOI:** 10.1101/2020.04.13.040170

**Authors:** Chee Wah Yuen, Mardani Abdul Halim, Nazalan Najimudin, Ghows Azzam

**Affiliations:** School of Biological Sciences, Universiti Sains Malaysia, 11800 Penang, Malaysia; USM-RIKEN International Centre for Ageing Science (URICAS), Universiti Sains Malaysia, 11800 Penang, Malaysia

**Author notes:** Corresponding authors Email address School of Biological Sciences, Universiti Sains Malaysia, 11800 Penang, Email address School of Biological Sciences, Universiti Sains Malaysia, 11800 Penang.

**Keywords:** Salvianolic acid A, Alzheimer’s disease, β-amyloid, *C. elegans*

## Abstract

Alzheimer’s disease (AD) is a brain disease attributed to the accumulation of extracellular senile plaques comprising β-amyloid peptide (Aβ). In this study, a transgenic *Caenorhabditis elegans* containing the human beta amyloid Aβ_42_ gene which exhibited paralysis when expressed, was used to study the anti-paralysis effect of salvianolic acid A. Various concentrations ranging from 1 μg/ml to 100 μg/ml of salvianolic acid A were tested and exhibited the highest effect on the worm at the concentration of 100 μg/ml. For anti-aggregation effect, 14 μg/ml salvianolic acid A (within 4 mg/ml of Danshen) showed a significant level of inhibition of the formation of Aβ fibrils. An amount of 100 μg/ml of salvianolic acid A had the potential in reducing the ROS but did not totally obliterate the ROS production in the worms. Salvianolic acid A was found to delay the paralysis of the transgenic *C. elegans*, decrease Aβ_42_ aggregation and decreased Aβ-induced oxidative stress.

## 1. INTRODUCTION

Alzheimer’s disease (AD) is a chronic, irrevocable and progressive brain disorder which affects middle or old age population leading to memory loss and thinking disability which subsequently resulted in interference in conducting simplest tasks in daily life. AD is believed to be explainable using the amyloid cascade hypothesis. The proposed amyloid hypothesis was supported with several evidences. The first evidence was the discovery of senile plaques (SPs) and neurofibrillary tangles (NFTs) by a physician, Dr Aloi Alzheimer, when autopsy was performed on an AD patient’s brain in 1907 (reviewed by Walsh and Teplow, 2012) [1]. This was followed by significant findings of beta-amyloid (Aβ) within SPs [2], discovery of <u>a</u>myloid <u>p</u>recursor <u>p</u>rotein (APP) gene sequence [3] mutations in it [4] which led to the proposed amyloid cascade hypothesis [5]. Briefly, the amyloid cascade hypothesis encompasses the scission of APP by β-secretase and γ-secretase to form Aβ peptides, followed by aggregation of Aβ oligomers to produce SPs and eventually causing toxicity to the brain cells by oxidative stress.

Based on the amyloid cascade hypothesis, a search for potential therapeutics that inhibits the Aβ production had been initiated. Among the potential therapeutics that hinders Aβ production are drugs that contain anti-Aβ aggregation properties to disrupt the formation of SPs and antioxidant effects to decrease the oxidative stress caused by Aβ. Since there is no disease modifying drugs for AD, the current drugs that are available can only delay the onset of AD [6]. However, there are drawbacks for these drugs in the effort to modify the AD pathogenesis and the deleterious side effects of the present drug, thus, there is a need to search for new potential disease modifying AD drugs.

Salvianolic acid A is one of the major water-soluble compounds in the water Danshen extract besides salvianolic acid B and danshensu [7]. Salvianolic acid A have exhibited its antioxidant properties by inhibiting ROS production due to H_2_O_2_ induction, and significantly scavenging HO‧ that was produced in phorbol myristate acetate-stimulated rat neutrophils [8, 9]. Studies had demonstrated that salvianolic acid A has the ability to improve memory impairment when the compound was injected intravenously in mice [10]. It was also reported that 10 mg/kg salvianolic acid A that was intravenously injected into the mice can inhibit cerebral lipid peroxidation and clear free HO‧ radicals. Hence, it can be deduced that there is a relationship between its antioxidant properties with its improving effects on memory impairment induced by cerebral ischemia-reperfusion in mice [10].

In this study, we used transgenic *Caenorhabditis elegans* carrying Aβ_42_ gene to investigate the potential effect of salvianolic acid A towards AD. Here we showed that salvianolic acid A have the ability to delay the paralysis of the worms and also reduce the ROS.

## 2. MATERIALS AND METHODS

### 2.1 *C. elegans* Strains and Maintenance

Transgenic *C. elegans* strain GMC101(dvIs100[unc-54p::A-beta-1-42::unc-54 3’-UTR + mtl-2p::GFP), *C. elegans* strain CL2122 (dvIs15 [(pPD30.38) unc-54(vector) + (pCL26) mtl-2::GFP] and *E. coli* OP50 strain was kindly provided by the Caenorhabditis Genetics Center (CGC), University of Minnesota (Minneapolis, MN, USA). The transgenic C. elegans were maintained at 16 °C on nematode growth medium (NGM) seeded with *E. coli* OP50 bacteria. GMC101 is a transgenic strain that expressed the human Aβ_42_ gene while CL2122 does not have the human Aβ_42_ gene.

### 2.2 *C. elegans* paralysis assays

The rapid paralysis phenomenon due to the expression of Aβ_42_ in *C. elegans* GMC101 strain was suitable to be exploited to screen for natural compounds with neuroprotective properties. Paralysis assays were performed as described by McColl *et al*. [11] with slight modifications by letting the adult worms to lay eggs for 4 hours and shifting the worms from 16°C to 25°C after 72 hours of post-egg laying. All populations were cultured at 16°C on 60 mm NGM plates with 250 μl of live *E. coli* OP50 bacterial culture pre-spread on the plates. The *C. elegans* populations were developmentally synchronized from a 4 hour egg-lay on NGM plates in the absence or presence of the drugs. After 72 hours of post egg-laying, the individuals were shifted to 25°C. Time zero was defined at this point of temperature shift. The body movement of the nematodes were assessed over time by scoring as “paralysed” if they failed to show complete full body movement, either spontaneously or touch-provoked. Proportion of individuals that were not paralysed was calculated. In all studies, three independent experiments were performed. A range of different concentrations of salvianolic acid A (1-100μg/ml) was used to feed the nematodes. Salvianolic acid A, was purchased from a Chinese manufacturer (Phytomarker, Tianjin, China). The purity of salvianolic acid A was all above 98%.

### 2.3 Measurement of reactive oxygen species (ROS) in *C. elegans*

Levels of ROS were measured *in vivo* using the 2,7-dichlorofuorescein diacetate method as described by Gutierrez-Zepeda *et al.* [12] with modifications by incubating the worms at 37°C between reads to simulate human body temperature and readings were taken every 20 minutes. The compound 2,7-dichlorofuorescein diacetate is able to permeate the cells of *C. elegans* and is intracellularly converted to highly fluorescent 2′,7′-dichlorofluorescein (H2DCFs) when it is oxidized. The worms were synchronized by performing the egg-laying process on NGM plates either containing no compound as a control, or containing salvianolic acid A at the desired concentrations. These were incubated at 16°C for 72 hrs. To initiate amyloid induced paralysis, the live worms were shifted to 25°C. Approximately 100 worms were harvested at 32 hrs after temperature shift using M9 buffer. The worms were resuspended in M9 buffer with 100 μM DCFH-DA (Sigma-Aldrich, Missouri, USA). Worms were transferred into the wells of 96 well non-binding black microplate (Greiner Bio-One) and read every 20 minutes at an excitation wavelength of 485 nm and an emission wavelength of 520 nm using a flourescence microplate reader (EnVision 2104 Multilabel Reader (Perkin Elmer, MA, USA). Worms in M9 buffer absent of DCFH-DA were used as negative controls. Nematodes were fed with different concentrations salvianolic acid A (1-100μg/ml).

### 2.4 Immuno-dot blot assay of Aβ

The levels of Aβ proteins were analysed using the immuno-dot blot analysis as described by Sola *et al*. [13] with modifications. Total proteins were visualized using colorimetric method in this study as opposed to the usage of Ponceau red solution as reported by Sola *et al*. [13]. For comparison of Aβ_42_ levels, approximately 1000 adults GMC101 strain that were treated or untreated with 100 μg/ml salvianolic acid A (Phytomarker) were collected in S-basal medium in three biological replicates. Samples were then extracted in 3 volumes of urea buffer (7 M urea, 2 M Thiourea, 4% w/v CHAPS, 1.5% w/v dithiothreitol and 50 mM Tris pH 8.0) disrupted via sonication, and then centrifuged at 13,000 x*g* for 10 min. Total proteins were spotted (10 μg) onto nitrocellulose membranes (Merck Millipore, Darmstadt, Germany) and left to dry. The membranes were incubated with a blocking solution [skim milk (Sunlac) with 0.1% tween-20-phosphate buffered saline] for one hour. This was followed by an incubation period with Anti-Aβ mouse monoclonal antibody 6E10 (1:1000, epitope: Aβ_3-8,_ Biolegend, San Diego) and peroxidase-conjugated anti-mouse IgG (1:1000, Bio-Rad, California), which acted as the primary and secondary antibodies, respectively. The spots were viewed using the colorimetric method available in Opti4CN^™^ Substrate Kit (Bio-Rad, California). Total protein extracted from CL2122 nematode strain was used as a negative control. The total protein extracted was quantitated using Bradford Reagent (Sigma, USA). Bovine Serum Albumin (BSA) (Sigma, USA) was used as the standard.

### 2.5 Aβ Aggregation assay

The procedure was performed as described by the manufacturer’s instructions (SensoLyte ® Thioflavin T Beta Amyloid (1-42) Aggregation Kit (Anaspec)). Thioflavin T is a dye that binds to the aggregated Aβ. Aggregation of Aβ was measured using 96 well non-binding black microplate (Greiner Bio-One). Approximately 10 μl of 2mM ThT, 85μL of Aβ_42_ peptide and 5μL of the bioactive compounds at selected concentrations were tested. The plates were read every 10 minutes on a fluorescence microplate reader (EnVision 2104 Multilabel Reader (Perkin Elmer, MA, USA) at an excitation wavelength of 440 nm with an emission wavelength of 484 nm. Salvianolic acid A at different concentrations was tested for their anti-Aβ aggregation effects. Morin (Anaspec) that was provided in the kit acted as a positive inhibitor.

## 3. RESULTS

### Salvianolic acid A delayed paralysis of the nematodes

Various concentrations of salvianolic acid A ranging from 1 μg/ml to 100 μg/ml were investigated to determine dose dependent effect towards the transgenic nematode strain GMC101. As shown in Fig. 1A and Fig. 1B, salvianolic acid A exhibited the highest effect on the worm at the concentration of 100 μg/ml compared to the untreated worms (p<0.05). No sign of paralysis was observed at the 32^nd^ hour of the experiment for concentration 100 μg/ml compared to the control which already showed the onset of paralysis. This indicated that the concentration of salvianolic acid A was effective in protecting against the toxicity produced by Aβ_42_. Even for the concentrations of 75 μg/ml (p<0.05) and 50 μg/ml (p<0.05) showed significant effects when compared to the control. However, there was no significant difference among the concentrations of salvianolic acid A ranging from 1 μg/ml to 12.5 μg/ml (p>0.05).

**Figure 1:**
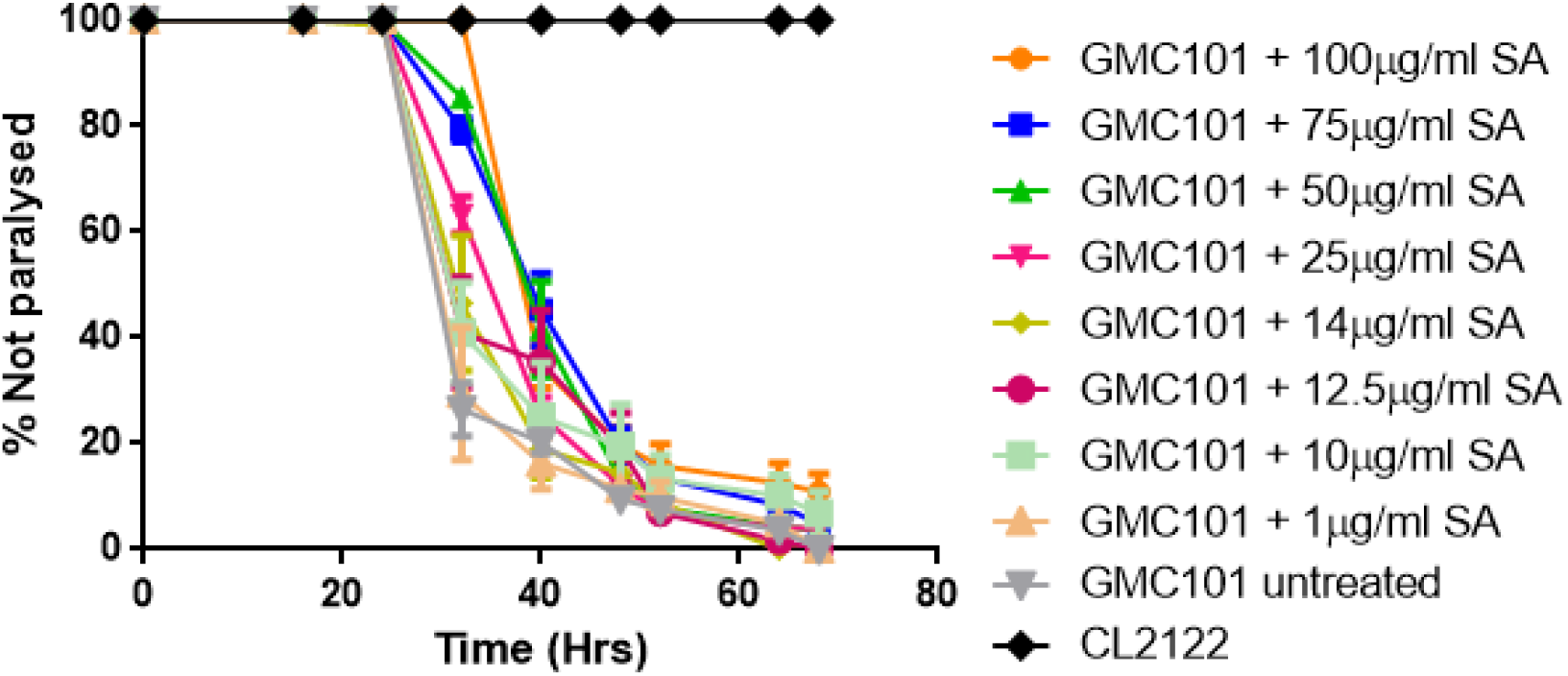
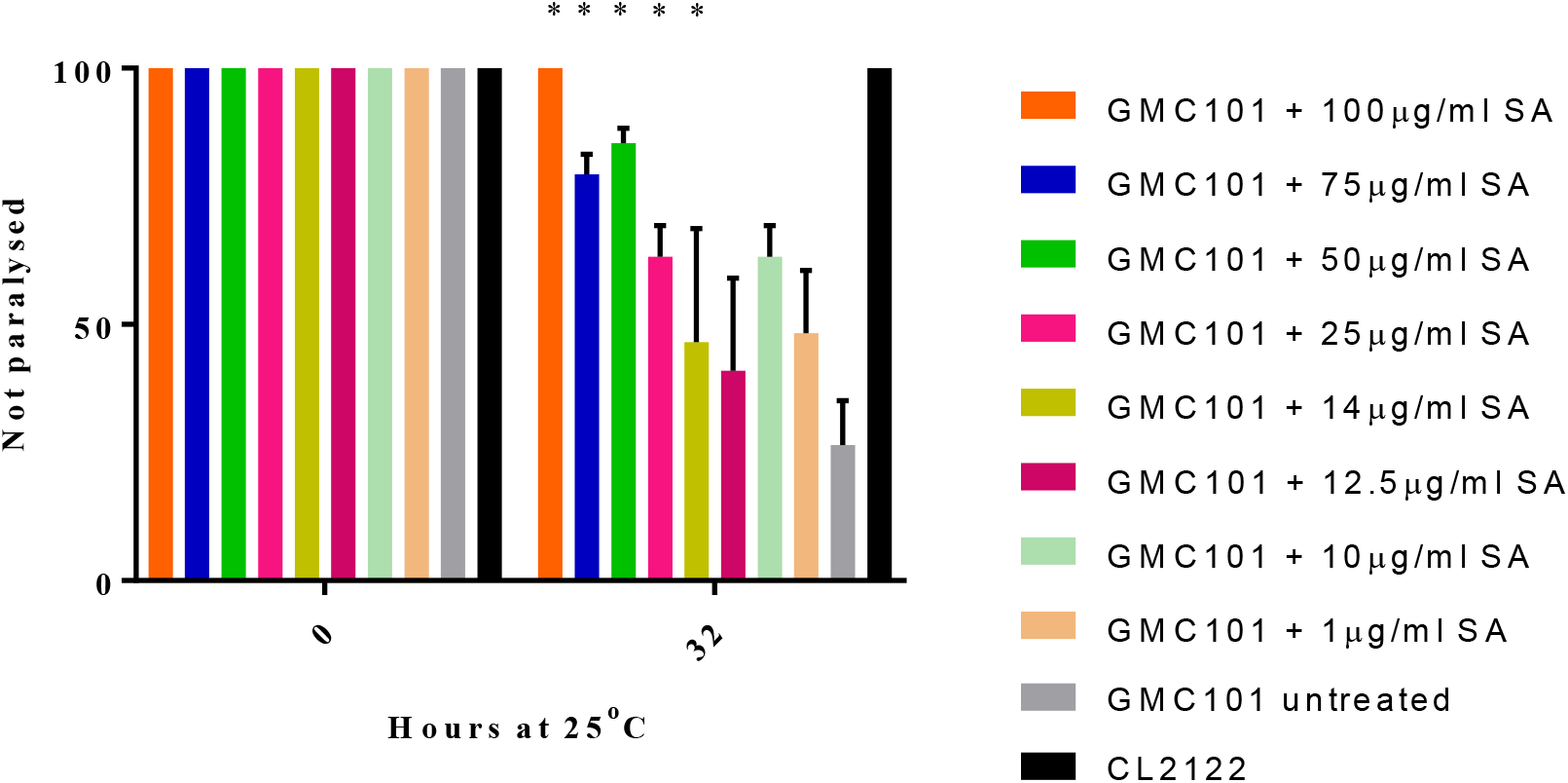
Effects of salvianolic acid A (SA) on the paralysis of the worms. **(A)** salvianolic acid A delayed Aβ induced paralysis in transgenic *C. elegans,* strain GMC101 at concentrations 1 μg/ml to 100 μg/ml salvianolic acid A. **(B)** Comparison of transgenic *C. elegans* strain GMC101 treated at various concentrations of salvianolic acid A at 0 and 32 hours of the paralysis assay. 50, 75 and 100 μg/ml salvianolic acid A significantly delayed paralysis of the GMC101 strain (p<0.05). Paralysis was also delayed when worms exposed to 25 μg/ml (p<0.05) and 14 μg/ml (p<0.05) salvianolic acid A at 32^nd^ hour after upshift from 16 °C to 25 °C.

#### 3.1 Salvianolic acid A inhibits Aβ aggregation

To determine whether salvianolic acid A had an anti-aggregation effect by inhibiting the formation of Aβ_42_ fibrils which was believed to be toxic to the demented brain, an *in vitro* aggregation assay using thioflavin T dye to measure Aβ_42_ aggregation was employed. A concentration of 4mg/ml Danshen water extract contained approximately 14 μg/ml of salvianolic acid A was used. A significant level of inhibition of the Aβ fibrils formation was obtained when 14 μg/ml salvianolic acid A was used compared to the other two lower concentrations of 7 μg/ml and 3.5 μg/ml (Fig. 2A). The inhibitory effect produced by 14 μg/ml salvianolic acid A was not as great as the positive control. The lowest concentration of 1.75 μg/ml salvianolic acid A tested did not stop Aβ aggregation (Fig. 2B). The IC_50_ for the inhibition of Aβ aggregation by salvianolic acid A obtained was 7 μg/ml.

**Figure 2:**
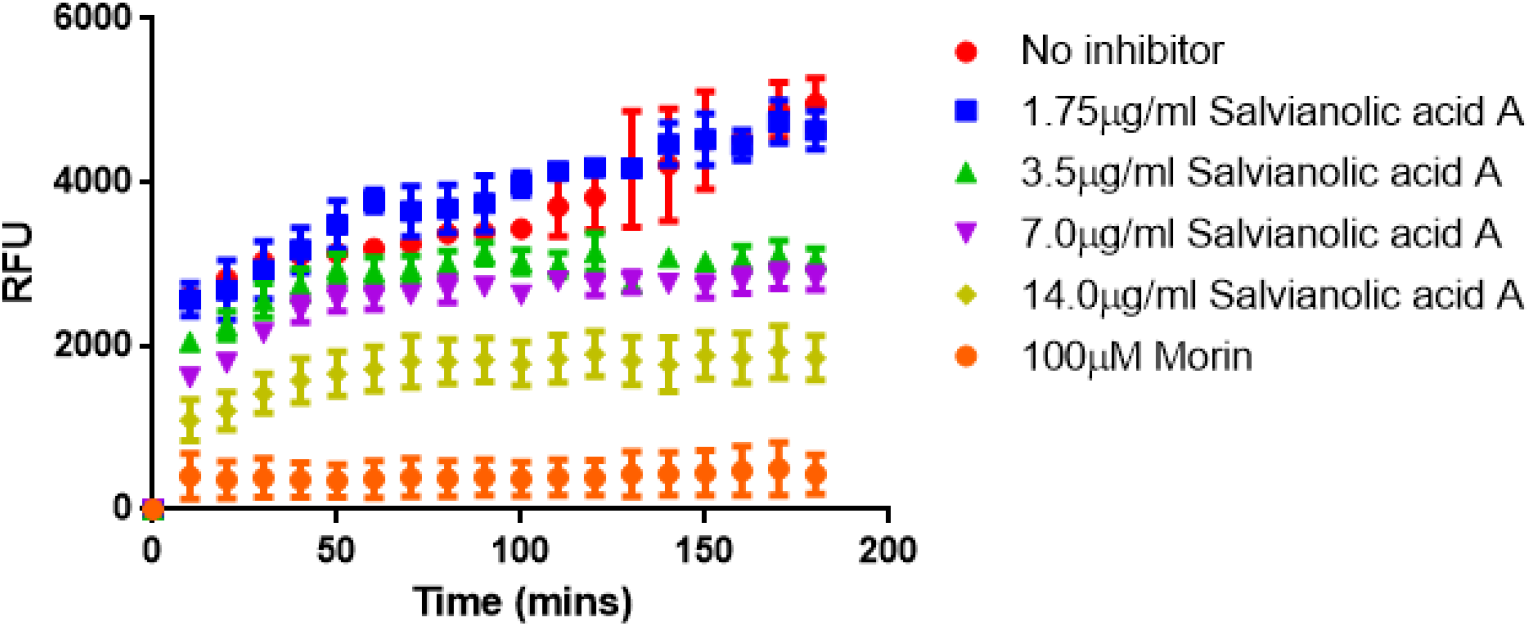
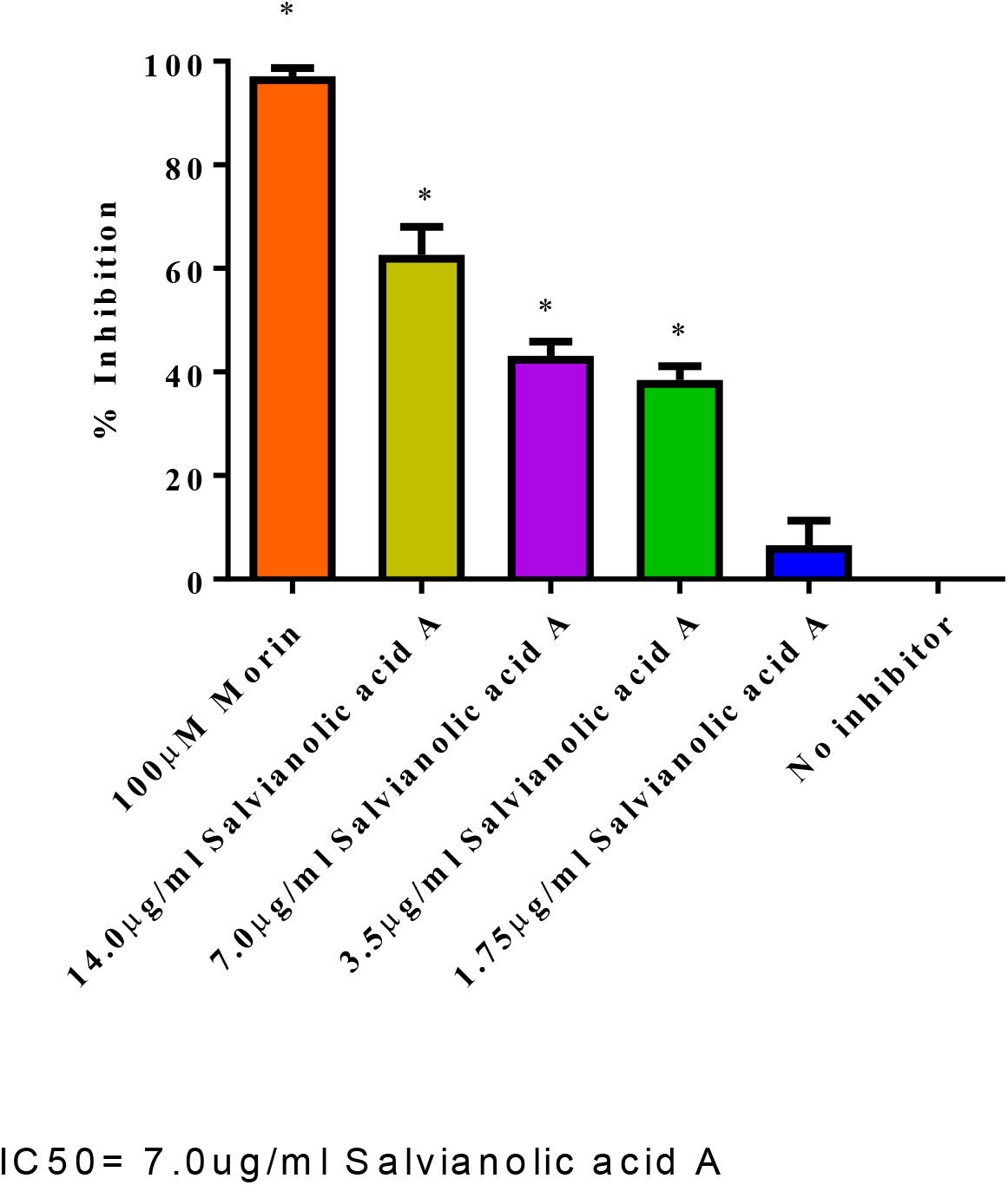
Effects of salvianolic acid A on Aβ aggregation. **(A)** Salvianolic acid A inhibits the Aβ_42_ formation comparing various concentrations of 1.75 μg/ml to 14μg/ml salvianolic acid A at a duration of 180 minutes. Fluorescence signal was monitored at Ex/Em= 440/484 nm every 10 minutes at 37 °C with 15 seconds between reads. 100μM morin acts as a positive inhibitor. **(B)** The percentage of inhibition of Aβ_42_ formation by various concentrations of salvianolic acid A at 180 minutes. Aβ_42_ aggregation was inhibited at concentrations of 3.5 μg/ml, 7.0 μg/ml (p<0.05) and 14.0μg/ml (p<0.05) salvianolic acid A as compared to the control. IC_50_ was 7.0 μg/ml salvianolic acid A.

#### 3.2 Effect of salvianolic acid A towards oxidative stress

Salvianolic acid A also had the potential of reducing ROS level (Fig 3A). Differences in ROS production were observed between worms treated and untreated with salvianolic acid A. As shown in Fig. 3(B), salvianolic acid A had the potential in reducing the ROS production but did not totally obliterate the ROS production. The results were in correlation with the anti-paralysis assays. The total Aβ protein that was extracted from transgenic nematodes GMC101 strain that were exposed to 100 μg/ml salvianolic acid A was not significantly reduced. This indicated that there was no decrease in the level of Aβ_42_ protein as can be seen in the dot blot experiment (p>0.05) (Fig 4A and Fig. 4B). No unspecific binding of Aβ antibody when total protein of CL2122 was used. Thus, based on the results obtained, it could be deduced that salvianolic acid A had both anti-aggregation and anti-oxidant properties. However, its effect on the total Aβ production was not significant.

**Figure 3:**
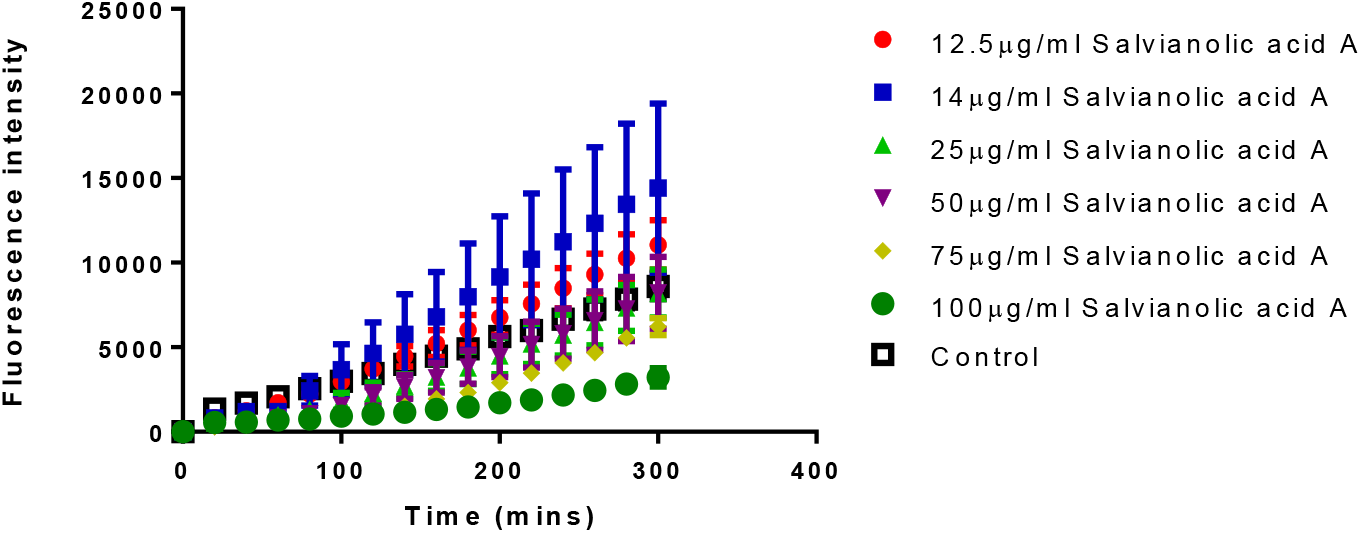
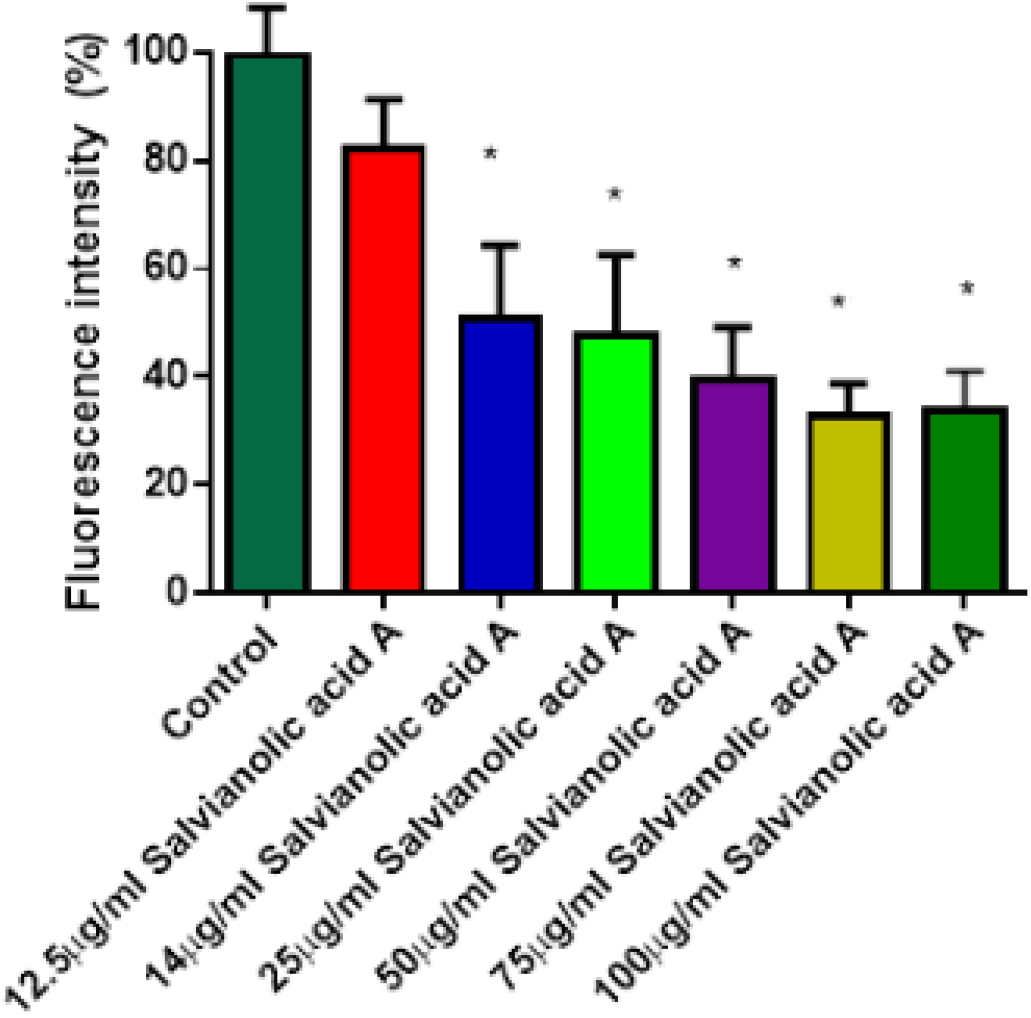
Effects of salvianolic acid A on the reduction of oxidative stress. **(A)** Comparison of relative oxygen species levels in transgenic *C. elegans* GMC101 treated with concentrations 12.5 μg/ml to 100μg/ml salvianolic acid A with the duration of 300 minutes with every 20 minutes readings. **(B)** Comparison relative oxygen species levels in transgenic *C. elegans* GMC101 treated with concentrations 12.5 μg/ml to 100 μg/ml at the 60 minutes. Oxidative stress in worms was significantly reduced at concentrations 14 μg/ml, 25 μg/ml (p<0.05), 50 μg/ml, 75μg/ml and 100 μg/ml (p<0.05) salvianolic acid A as compared to the control.

**Figure 4:**
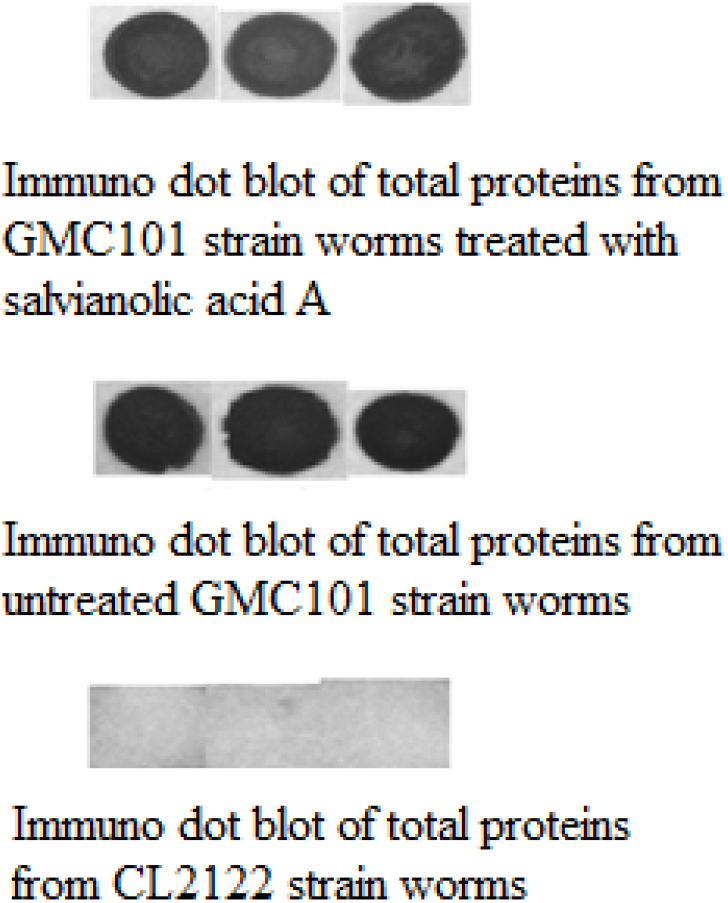
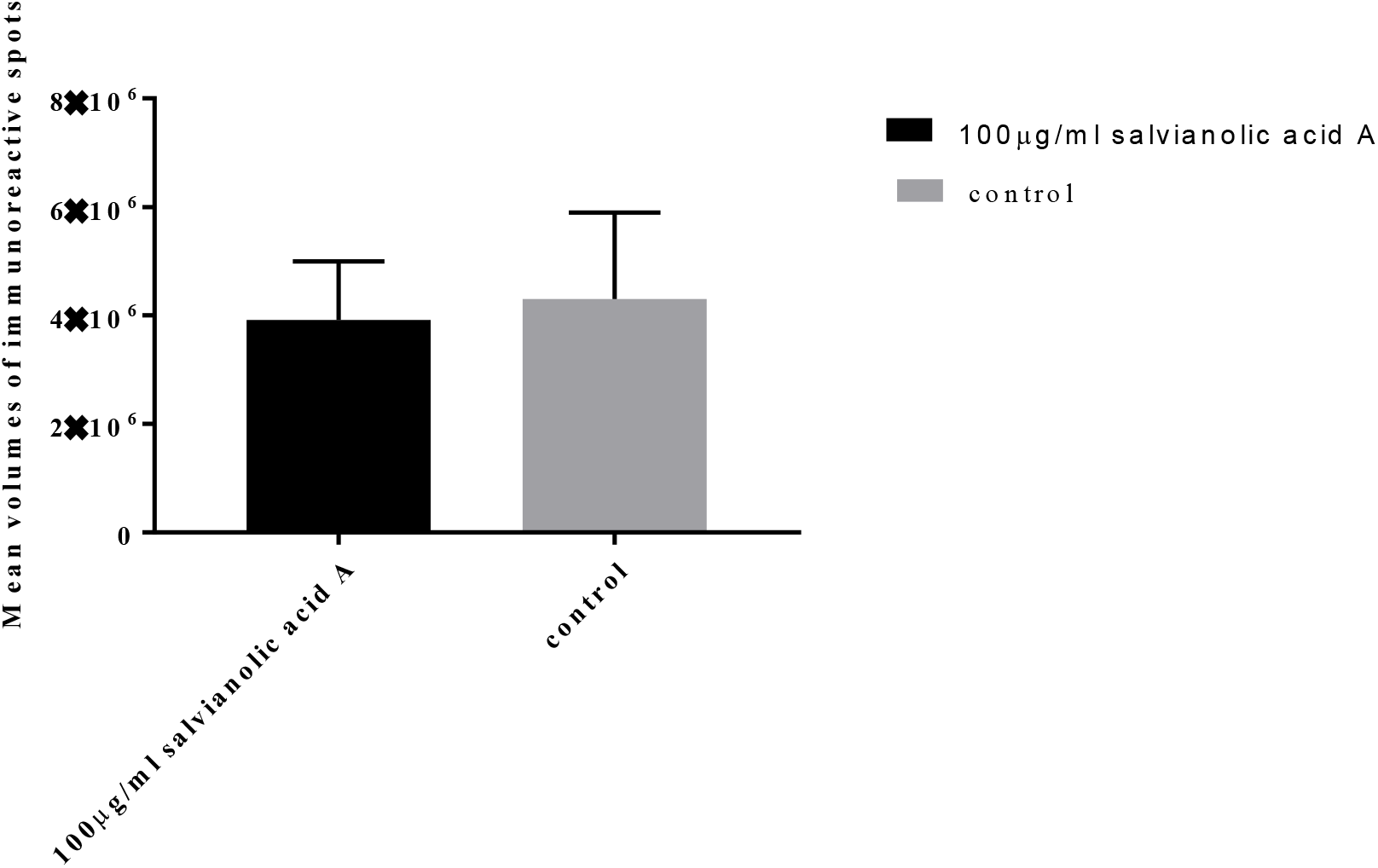
Effects of salvianolic acid A on the Aβ levels. **(A)** Representative dot blot of total Aβ in GMC101 transgenic worms fed with or without 100 μg/ml salvianolic acid A. No distinctive difference in intensity of dot blot results between the Aβ protein that were extracted from worms that fed with or without salvianolic acid A. No unspecific binding of Aβ antibody observed when total proteins of CL2122 were used. **(B)** Exposure to 100 μg/ml salvianolic acid A for 32 hours did not significantly alter total Aβ levels (p>0.05).

## 4. DISCUSSION

According to the amyloid cascade hypothesis, soluble Aβ oligomers aggregate to form protofibrils and subsequently assemble to produce insoluble Aβ fibrils and plaques. These fibrils are believed to be neurotoxic that cause the onset of AD [14–16]. The stability of the antiparallel β-sheet structure of Aβ is due to the aromatic interactions between Aβ peptides which subsequently leads to formation of Aβ fibrils [17–19]. Salvianolic acid A was tested for its potential anti-Aβ_42_ aggregation property based on several reports showing that polyphenols had potentially decrease aromatic interactions between Aβ peptides [20, 21]. Salvianolic acid A had been demonstrated to hinder the Aβ_42_ fibrillogenesis by blocking α-helices to form β-sheets and disaggregate preformed Aβ_42_ fibril [22]. Inhibitory effects of salvianolic acid A towards Aβ_42_ aggregation in this study is in agreement with the study reported. In a separate study, metal ions such as Cu (II), Fe (III), and Zn (II) were shown to induce Aβ_42_ aggregation and this phenomenon can be inhibited through salvianolic acid A chelation. In addition, the same study also shown that salvianolic acid A also inhibits Aβ_42_ self-mediated aggregation as well as disaggregated Aβ_42_ ageing fibrils [22]. Hence, it can be assumed that salvianolic acid A, contributed to the inhibitory effects of the extract.

Numerous reports have shown that salvianolic acid A has antioxidant properties. Salvianolic acid A disrupted H_2_O_2_-induced ROS production, and scavenges HO. in phorbol myristate acetate-stimulated rat neutrophils [8, 9]. Further studies had shown that salvianolic acid A reduced oxidative stress due to accumulation of ROS in SH-SY5Y [22]. In addition, salvianolic acid A inhibited cerebral lipid peroxidation and diminishes free HO. radicals which subsequently decreased memory impairment caused by cerebral ischemia-reperfusion in mice [10]. In another study, 20 mg/kg of salvianolic acid A significantly decreased hepatoxixity and also reduced oxidative stress in CCl_4_-induced rats by the observation of decreased in reactive oxygen species production and also malondialdehyde concentration in the liver tissues [23]. In comparison, the current study had shown that salvianolic acid A reduced oxidative stress that was accumulated in the transgenic *C. elegans* carrying Aβ_42_.

## 5. CONCLUSION

In this study, we found that salvianolic acid A treatment towards *C. elegans* expressing human Aβ_42_ gene showed encouraging response. From the study, paralysis was deferred when *C. elegans* was treated with salvianolic acid A. In addition, it also inhibits the development of Aβ fibrils as well as decreasing Aβ-induced ROS production in *C. elegans*. Through this study, we conclude that salvianolic acid A has the potential to be an alternative drug to combat Alzheimer’s disease.

## Acknowledgements

We would like to thank all our collaborators and colleagues for the discussion and the work conducted in this lab. This study was funded by the USM Top Down Research Fund - URICAS (1001/PBIOLOGI/870029).

